# Acetone-Butanol-Ethanol (ABE) fermentation with Clostridial Co-cultures for Enhanced Biobutanol Production

**DOI:** 10.1101/2023.12.08.570763

**Authors:** Karan Kumar, Shraddha M. Jadhav, Vijayanand S. Moholkar

## Abstract

This study investigates acetone-butanol-ethanol (ABE) fermentation, a process using solventogenic *Clostridium* species to produce acetone, butanol, and ethanol. Recent biotechnological advancements, such as omics, systems biology, and metabolic engineering, have reignited interest in butanol production, responding to the increasing gasoline costs and the demand for sustainable energy systems. This study unravels the distinct physiological attributes of *C. acetobutylicum* (*Cac*) and *C. pasteurianum* (*Cpa*), significantly impacting sustainable bioenergy technologies. Employing response surface methodology (RSM), we embarked on a comprehensive statistical optimization journey in the co-culture system, *Cac* MTCC 11274 and *Cpa* MTCC 116, enhancing biobutanol production from mixed substrates. A spectrum of process parameters was scrutinized, encompassing the ratio of *Cac* and *Cpa* inoculum, sodium concentration, and the ratio of xylose to glucose. Statistical analysis revealed salt concentration’s profound influence on biomass, total alcohol, and butanol production. The culmination of these endeavors yielded highly promising outcomes: a butanol concentration of 12.1 ± 0.45 g L^-1^ (model prediction: 11.87 g L^-1^), biomass of 4.15 ± 0.03 (model prediction: 4.06 OD_600_), and ABE concentration of 23.1 ± 0.55 g L^-1^ (model prediction: 22.45 g L^-1^). These results represent a significant leap forward in bioenergy technologies, offering both practical insights and sustainable solutions for enhanced biofuel production.

## 1. Introduction

Alcoholic biofuels (such as bioethanol and biobutanol) have emerged as potential alternate and renewable liquid transportation fuels (Kumar et al., 2021, 2023; Kumar and Moholkar, 2023). Among these, biobutanol holds special potential and promise due to its similar properties as gasoline (air:fuel ratio = 12.2, energy density = 29.2 MJ/L, research octane number = 96) (Kuszewski, 2018; Yogesh, 2022). Butanol blends with gasoline up to 80% v/v are compatible with the current internal combustion engines in vehicles (Kolesinska et al., 2019; Haifeng Liu et al., 2019). Acetone-Butanol-Ethanol (ABE) fermentation process using solventogenic clostridia for production of butanol is known for more than a century (Kumar et al., 2023). ABE fermentation was a major industrial process for production of acetone and butanol till mid-1960s, but it was almost extinct with fast development and growth of the petrochemical industry after WW II (Jones et al., 2023; Li et al., 2020). However, in recent years, butanol production through biological routes has seen renewed interest of the academic and industrial fraternity due to two reasons: (1) climate change risk and serious global energy security issue, which have triggered extensive research in carbon neutral fuels from renewable and sustainable sources (Chang et al., 2022; Kolesinska et al., 2019), and (2) advances in microbiology and biotechnology that have resulted in more efficient mutants and genetically modified microbial strain with higher yield, solvent tolerance, and selectivity for butanol (Boudignon et al., 2023; Bourgade et al., 2021; Cheng et al., 2019).

Lignocellulosic biomass (LB) in the form of agro- and forest residues, available abundantly throughout the year, is potential sustainable feedstock for fermentation (Birgen et al., 2019; Costa et al., 2020). Two main components of LB, viz. cellulose and hemicellulose, can be hydrolysed to produce hexose-rich and pentose-rich hydrolysates which can be fermented by clostridial cultures to acetone, butanol, and ethanol (Costa et al., 2020; Kucharska et al., 2018; Liu et al., 2021).

Most of the previous literature has employed as single clostridial strain for ABE fermentation. More recently, co-cultures of clostridia have also been used in ABE fermentation due to their distinct merits (Cui et al., 2021; Du et al., 2020). Either two solventogenic clostridia or solventogenic + cellulolytic/gas-fermenting clostridia have been coupled in the co-culture fermentation. The major merits of clostridial co-culture systems are as follows: (1) effective utilization of a wider range of complex substrates (especially the lignocellulosic biomass from different sources), (2) improvement in product yield and product rates, (3) more robust fermentation system with higher tolerance to O_2_ and solvents, (4) widening of product synthesis chain with utilization of main substrates (e.g. carbohydrates), and also by-products (e.g. CO and CO_2_) that results in higher product yields (Cui et al., 2021; Gebreselassie and Antoniewicz, 2015). The co-culture system has also been designated as “Consolidated Bioprocessing (CBP)”, which comprises the individual steps of in-situ secretion of saccharolytic enzymes by one of the bacterial cultures, hydrolysis of cellulose/ hemicellulose by the saccharolytic enzymes, and fermentation of monomeric sugars by one or both bacterial cultures (Du et al., 2020; Jiang et al., 2020; Wen et al., 2020).

In the present study, we have employed the co-culture of two solventogenic clostridia, viz. *C. acetobutylicum* and *C. pasteurianum*, for fermentation of mixed substrates comprising pentose (xylose) and hexose (glucose) sugars for enhanced butanol production. Both of these clostridial strains have been extensively investigated for ABE fermentation with different carbon sources (Diallo et al., 2021; Li et al., 2020). We conjectured that combining these two strains would result in better utilization of different carbon sources in the hydrolysates from lignocellulosic biomass. Initially, the total solvent (ABE) production and butanol selectivity of individual clostridial cultures grown on different pentose and hexose sugars found in LB hydrolysates was assessed. Thereafter, the CBP of ABE fermentation with co-culture system was optimized using the statistical design of experiments. The synergistic microbial interactions among the two clostridial cultures have resulted in enhanced butanol production.

## 2. Material and methods

### 2.1. Bacterial strains, maintenance, and inoculum preparation

All chemicals were obtained from HiMedia Pvt. Ltd., India, and were used in their original form. Cultures of *Cac* and *Cpa* with identification numbers MTCC 11274 and MTCC 116, respectively, were sourced from the Microbial Type Culture Collection (MTCC) in Chandigarh, India. The lyophilized cultures of *Cac* and *Cpa* were individually reanimated on Reinforced Clostridial Agar (RCA) plates. The plates were then transferred to a vacuum desiccator, where anaerobic conditions were maintained using an Anaerogas pack (LE002A, HiMedia). Afterward, the cultures were placed in an incubator at 37 °C. In 1 liter of distilled water, the composition of RCA was as follows: peptone 10.0 g, yeast extract 5.0 g, glucose 5.0 g, beef extract 10.0 g, starch 1.0 g, NaCl 5.0 g, L-cysteine HCl 0.5 g, CH_3_COONa 3.0 g, and agar 0.5 g. The medium’s pH was set to 6.8 ± 0.2 by incorporating 1 M NaOH. To commence the inoculation procedure, a 120 mL serum bottle was utilized, containing 50 mL of RCM medium (RCA without agar). This setup was positioned within a rotary incubator shaker (Manufacturer: Lab Companion; Model: SI–300R) and subjected to incubation at 37 LJ with a rotational speed of 150 rpm for a duration of 24 hours. The resulting culture broth served as stock and was subcultured monthly. Overnight-grown cultures of both clostridial strains were individually utilized as inoculants for batch fermentation experiments.

### 2.2. Batch fermentation with glucose as substrate (base case)

Batch trials with separate cultures commenced by transferring the inoculum from the mid-log phase of the culture into a 120 mL serum bottle. This bottle contained 60 mL MoBP medium, a modified medium optimized for butanol productivity, as detailed by Ahlawat et al., (2019). The total 60 mL quantity comprised of 52.8 mL of the MoBP medium, 1.2 mL (3% w v^-1^) L–cysteine HCl, and 6 mL (10% v v^-1^) of *Clostridium* inoculum. Additionally, 0.1% w v^-1^ Resazurin dye was incorporated into the medium to indicate anaerobic conditions. The MoBP medium consisted of the following components with specified concentrations (g L^-1^): peptone (49.7), glucose (80.0), KH_2_PO_4_ (0.5), K_2_HPO_4_ (0.5), FeSO_4_·7H_2_O (0.023), MnSO_4_·H_2_O (0.023), NaCl (5), MgSO_4_·7H_2_O (0.46), CH_3_COONH_4_^+^ (2.2), biotin (0.01). Additionally, 10 mL each of trace and vitamin solution were added to 1 L of MoBP medium. The trace element solution consisted of the following components with specified concentrations (in g L^-1^): H_3_BO_3_ (0.01), Na_2_MoO_4_ (0.01), N(CH_3_COO)_3_ (4.5), AlK(SO_4_)_2_ (0.01), CaCl_2_·2H_2_O (0.01), ZnSO_4_·7H_2_O (0.05), CoCl_2_·6H_2_O (0.2), CuCl_2_·6H_2_O (0.05). The composition of the vitamin solution was as follows (concentration in g L^-1^): folic acid (0.01), riboflavin (0.025), para–amino benzoic acid (0.01), and citric acid (0.02). Before introducing the inoculum, the medium underwent a 15-minute purge with pure nitrogen (99.99%). Following that, the system was sterilized at 121 °C and 15 psi for 15 minutes in autoclave. Afterwards, the bottles were inoculated with a 10% v v^-1^ inoculum of *Cac* and *Cpa* individual cultures, then incubated at 37 °C and 150 rpm. To ensure result reproducibility, all experiments were replicated in triplicate. Periodic sampling of fermentation broth aliquots was conducted to quantify the glucose consumption over time.

### 2.3. Assessment of growth of *Cac* MTCC 11274 and *Cpa* MTCC 116 in alternate carbon sources

As noted previously, the hydrolyzates obtained from lignocellulosic biomass (after purification and removal of inhibitors) contain numerous pentose and hexose sugars, in addition to other compounds like sugar alcohols and polysaccharides that can be potential carbon sources. Enhancement of fermentation yield would necessitate effective utilization of all components present in the hydrolysates by the microbial culture. Therefore, in the initial phase of our study, we conducted screenings for the growth profiles and butanol production of each of the two clostridial strains. This involved utilizing a total of 15 different carbon sources, encompassing three three disaccharides (maltose, sucrose, and lactose), three polysaccharides (starch, dextrin, and cellulose), four hexoses (fructose, galactose, mannose, and dextrose), pentoses (arabinose, ribose, and xylose), and two sugar alcohols (glycerol and mannitol). The experimental approach involved substituting glucose with each alternate carbon source in the composite MoBP medium, equivalent to 2.66 M of total carbon, while keeping all other components constant. To prevent any potential transfer of residual glucose from the seed medium to the test medium (with an alternate carbon source) during inoculation, the inoculum was subjected to centrifugation at 5,000×g for 10 minutes at 4 °C before inoculation and then re-suspended in the respective media. The sampling frequency and experimental conditions were maintained consistently, as detailed in section 2.2. The cell density and butanol concentration in the samples were assessed for all the carbon sources as substrates.

### 2.4. Optimization of co-culture parameters using statistical experimental design

Previous authors have shown that ABE fermentation in co-culture system grown over lignocellulosic hydrolysate is mainly influenced by the relative cell densities (or populations) of the two cultures (Barca et al., 2016; Muhammad et al., 2023), and relative concentrations of hexose and pentose carbon sources (Du et al., 2020). In addition to these parameters, the performance of the ABE fermentation system is also reported to depend on the sodium concentration in hydrolysates (Zhao et al., 2016). A quantitative metabolomics analysis by Zhao et al. (2016) has shown that high sodium concentrations affected the acidogenesis phase of cell metabolism with accumulation of NADH and ATP, and inhibition of central carbon metabolic (viz. glycolytic and pentose-phosphate) pathways. However, the ratio of NADP^+^/NADPH remained constant during fermentation leading to higher solvent specific productivities. The overall influence of these simultaneous phenomena was inhibition of biomass growth with greater solvent tolerance and higher solvent productivity. In view of these interesting results, we have also included the sodium concentration in the composite lignocellulosic hydrolysate medium as an optimization parameter.

Guided by these hypotheses, the optimization of ABE fermentation process parameters was conducted using response surface methodology (RSM) with a 3-factor Box-Behnken Design (BBD) of experiments. Three independent variables, namely inoculum ratio (Cac/Cpa), sodium concentration, and substrates ratio (xylose/glucose), were selected, while biomass, butanol, and ABE titers (or concentrations) were designated as the response variables. These parameters were investigated across three levels, as outlined in Table 1A, and the ranges were determined based on the outcomes of the initial batch experiments.

**Table 1(A):** Factors and levels of optimization variables in Box-Behnken Design.

| Variables | Coded values (Actual values) |  |
| --- | --- | --- |
|  | Upper bound | Lower bound |
| (X<sub>1</sub>) – Inoculum ratio (*Cac/Cpa*) | –1 (0.3) | +1 (1) |
| (X<sub>2</sub>) - NaCl (g L<sup>-1</sup>) | –1 (0) | +1 (10) |
| (X<sub>3</sub>) – Substrate ratio (Xyl/Glc) | –1 (0) | +1 (0.3) |

#### 2.4.1. Box-Behnken Design (BBD)

The BBD is a design approach widely employed in response surface methodology, serving as a tool to validate a 2^nd^order polynomial. It is commonly utilized for studying interactive effects among process variables. The experimental set was generated using Minitab statistical software (Release 16, Trial Version) with three coded levels (−1, 0, +1), three center points, and one replicate, as specified in Table 1A. The BBD generated runs are calculated using the following mathematical expression:

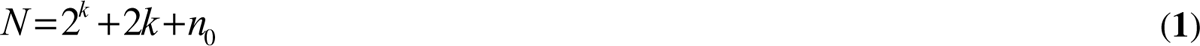

Here, N represents the number of runs, k is the number of independent parameters, and n_O_ represents the number of replicates runs at the center point. In this study, the experimental design consisted of 15 individual runs. The average response variables, including concentrations of biomass, butanol, and ABE, were recorded for each run, which was carried out in triplicates. The analysis of the individual and interactive effects of parameters on the response variable was performed using the following second-order quadratic model, as presented in Table 1B.

**Table 1(B):** Experimental and model predicted values of Box-Behnken Design for various response variables in co-culture systems.

| Expt. No. | Inoculum ratio (X<sub>1</sub>) | NaCl (X<sub>2</sub>) | Substrate ratio (X<sub>3</sub>) | Butanol (g L<sup>-1</sup>)<br>(Experiments) | Butanol (g L<sup>-1</sup>)<br>(Model) | Biomass (OD<sub>600</sub>)<br>(Experiments) | Biomass (OD<sub>600</sub>)<br>(Model) | ABE (g L<sup>-1</sup>)<br>(Experiments) | ABE (g L<sup>-1</sup>)<br>(Model) |
| --- | --- | --- | --- | --- | --- | --- | --- | --- | --- |
| 1 | 0.30 | 5 | 0.00 | 11.90 ± 0.40 | 11.84 | 3.99 ± 0.06 | 3.99 | 22.40 ± 0.45 | 22.31 |
| 2 | 1.00 | 5 | 0.30 | 7.40 ± 1.15 | 7.46 | 3.86 ± 0.05 | 3.86 | 14.70 ± 0.29 | 14.79 |
| 3 | 0.30 | 0 | 0.15 | 5.60 ± 0.96 | 5.63 | 4.07 ± 0.08 | 4.05 | 9.90 ± 0.19 | 10.22 |
| 4 | 1.00 | 0 | 0.15 | 4.80 ± 0.53 | 4.79 | 4.01 ± 0.07 | 4.02 | 9.60 ± 0.19 | 9.61 |
| 5 | 1.00 | 5 | 0.00 | 11.20 ± 0.45 | 11.18 | 3.88 ± 0.04 | 3.85 | 21.90 ± 0.44 | 22.12 |
| 6 | 0.65 | 0 | 0.00 | 10.10 ± 0.29 | 10.13 | 3.79 ± 0.03 | 3.82 | 19.70 ± 0.39 | 19.47 |
| 7 | 0.30 | 10 | 0.15 | 5.20 ± 0.53 | 5.21 | 3.99 ± 0.06 | 3.97 | 10.00 ± 0.2 | 9.98 |
| 8 | 0.65 | 5 | 0.15 | 5.94 ± 1.23 | 5.93 | 3.82 ± 0.02 | 3.82 | 11.24 ± 0.22 | 11.75 |
| 9 | 1.00 | 10 | 0.15 | 4.98 ± 0.53 | 4.95 | 3.71 ± 0.06 | 3.74 | 9.96 ± 0.19 | 9.64 |
| 10 | 0.65 | 0 | 0.30 | 7.40 ± 1.03 | 7.35 | 3.81 ± 0.05 | 3.79 | 14.65 ± 0.29 | 14.55 |
| 11 | 0.30 | 5 | 0.30 | 7.89 ± 1.25 | 7.91 | 3.94 ± 0.07 | 3.98 | 15.78 ± 0.32 | 15.56 |
| 12 | 0.65 | 5 | 0.15 | 5.92 ± 1.04 | 5.93 | 3.82 ± 0.05 | 3.82 | 12.12 ± 0.24 | 11.75 |
| 13 | 0.65 | 5 | 0.15 | 5.94 ± 0.98 | 5.93 | 3.82 ± 0.04 | 3.82 | 11.88 ± 0.24 | 11.75 |
| 14 | 0.65 | 10 | 0.30 | 6.20 ± 0.49 | 6.17 | 3.66 ± 0.04 | 3.64 | 12.10 ± 0.25 | 12.33 |
| 15 | 0.65 | 10 | 0.00 | 10.98 ± 0.28 | 11.03 | 3.59 ± 0.03 | 3.61 | 21.38 ± 0.43 | 21.48 |

In this mathematical expression:

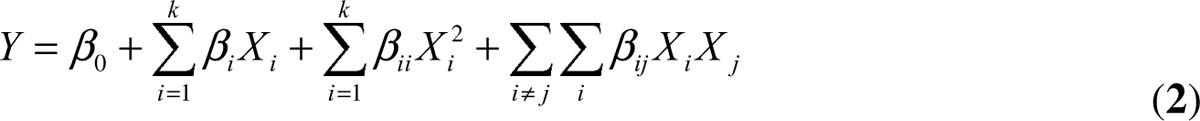

*Y* represents the measured response variable (concentrations of biomass, butanol, and ABE), β_0_ is the regression constant, *k* is the number of medium components or factors, β_i_ denotes the linear coefficient, β_ij_ signifies the interaction coefficient, β_ii_ represents the quadratic coefficient.

Experimental variables were transformed into coded values using Equation 3.

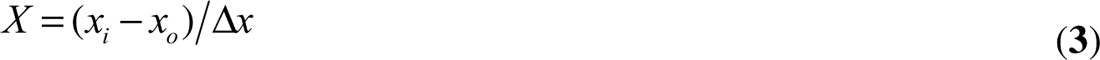

In Equation 3, the variables are defined as follows:

*X* is the dimensionless value of the variable, x_0_ is the variable value of x at the center point, Δ_x_ represents the step change.

The relative influence of the factor components on the response variable was assessed using Analysis of Variance (ANOVA). Interactive effects between independent variables (or factor components) were evaluated through response surface plots and contour plots.

### 2.5. Analytical methods

A volume of 1.5 mL from the fermentation culture was collected and centrifuged at 10,000×g for duration of 10 minutes at 4 °C. Afterwards, the pellet was washed and re-suspended in saline solution (0.85% w w^-1^ of NaCl). The measurement of growth was performed by assessing the optical density (OD) at 600 nm using a UV-visible spectrophotometer (Cary 50, Varian, Australia). Simultaneously, the supernatant was analyzed to generate dynamic profiles for organic acids (acetate and butyrate), pH, solvents (acetone, butanol, and ethanol), as well as residual substrate using high-performance liquid chromatography (HPLC, Model: Series 200, Manufacturer: Perkin Elmer, USA) equipped with an autosampler. Product separation was achieved through an Aminex-HPX 87H HPLC column (Model: Aminex HPX-87H, Manufacturer: BIO-RAD, USA) operating at 42 °C in column oven. Acetate and butyrate were detected using an UV detector at 210 nm, while solvent products and substrates were measured with a refractive index detector (RID). The estimated minimum detection limits for organic acids, solvents, and substrates were 0.04 g L^-1^, 0.06 g L^-1^, and 0.1g L^-1^, respectively. The correlation standard curves derived from the experiments are provided in the supplementary material. All measurements, representing the mean values along with their corresponding standard errors, were conducted in triplicate.

### 2.6. Validation experiments

Three additional fermentation test experiments were conducted at optimal parametric conditions to evaluate the accuracy of the optimal conditions predicted by the statistical analysis. The experiments were conducted in triplicate to ensure reproducibility of the results.

## 3. Results and discussion

### 3.1. Comparative growth characteristics of *Cac* MTCC 11274 and *Cpa* MTCC 116

#### 3.1.1. Comparative phenotypic/biochemical characterization

The comparative phenotypic or biochemical characterization of *Cac* MTCC 11274 and *Cpa* MTCC 116 are reported in Table 2. Both of the cells were motile and rod-shaped. Upon testing, both strains tested Negative for urease production, oxidase production, indole production, lipase production, nitrate production, and H_2_S production; however, they were positive for H_2_ gas production, which are typical biochemical characteristics of anaerobic *Cac* and *Cpa* strains.

**Table 2.** Phenotypic characteristics of *Cac* MTCC 11274 and *Cpa* MTCC 116.

| Phenotypic traits | *Cac* MTCC 11274 | *Cpa* MTCC 116 |
| --- | --- | --- |
| Cell morphology | Rods | Rods |
| Spore induction | Positive | Positive |
| Oxidase test | Negative | Negative |
| Motility | Positive | Positive |
| Indole production | Negative | Negative |
| Urease production | Negative | Negative |
| Lipase production | Negative | Negative |
| Nitrate reduction | Negative | Negative |
| H<sub>2</sub> production | Positive | Positive |
| H<sub>2</sub>S production | Negative | Negative |
| Optimum growth temperature | 37 | 37 |
| Optimal pH range | 4.8-6.8 | 5-7 |
| Optimum shaking | 150 – 160 rpm | 150 – 160 rpm |
| Major fermentation products | Organic acids, solvents, and gases (H<sub>2</sub>, and CO<sub>2</sub>) | Organic acids, solvents, and gases (H<sub>2</sub>, and CO<sub>2</sub>) |

#### 3.1.2. Comparative growth and fermentation characterization on glucose

Post-comparative biochemical characterization, both strains were compared for their growth and fermentation characteristics. We compared the strains’ growth and ABE production capabilities on MoBP media (Ahlawat et al., 2019; Kaushal et al., 2017). The rationale for choosing this media was that the strains’ maximum performance is very specific to the medium in which they are grown (Khamaiseh et al., 2014; Kumar et al., 2022; Lan et al., 2015).

The growth profiles and fermentation characteristics of *Cac* MTCC 11274 and *Cpa* MTCC 116 were investigated when cultured on glucose as the sole carbon source. The cultivation period spanned 120 hours, during which their growth and fermentation patterns were monitored. *Cpa* exhibited robust growth, reaching its highest optical density (OD_600_) of 2.45 at 28 hours, followed by a slight decline during the stationary phase. In contrast, *Cac* demonstrated a slower growth rate, with a maximum OD_600_ of 2.3 reached at 36 hours. This indicates that *Cac* has a slower glucose growth rate than *Cpa*. The fermentation characteristics were evaluated by measuring the production of acetate, butyrate, and butanol over the cultivation period. Both species exhibited typical ABE fermentation patterns.

*Cac* displayed efficient butanol production, reaching a maximum titer of 9.9 g L^-1^ at 40 hours. The butanol yield peaked at 0.21 g g^-1^ glucose, indicating a high efficiency in converting glucose to butanol. Acetate and butyrate were also produced, with maximum titers of 4.7 g L^-1^ and 0.95 g L^-1^, respectively, at 24 hours. On the other hand, *Cpa* showed lower butanol production compared to *Cac*. The maximum butanol titer obtained was 7.7 g L^-1^ at 36 hours, and the butanol yield reached 0.16 g g^-1^ glucose. The acetate and butyrate production by *Cpa* were slightly higher, with maximum titers of 5.1 g L^-1^ and 1.1 g L^-1^, respectively, at 24 hours. The time profile for biomass production, pH change, glucose consumption, solvents (acetone, ethanol, butanol, and total), and acids (acetate and butyrate) production are shown in Fig. 1. The differences in butanol production and fermentation characteristics between the two species highlight their distinct metabolic capabilities. *Cac* demonstrated superior butanol production and higher overall solvent (ABE) yields on glucose compared to *Cpa*. These findings suggest that *Cac* may be more suitable for biofuel production using glucose as a carbon source.

**Fig. 1.**
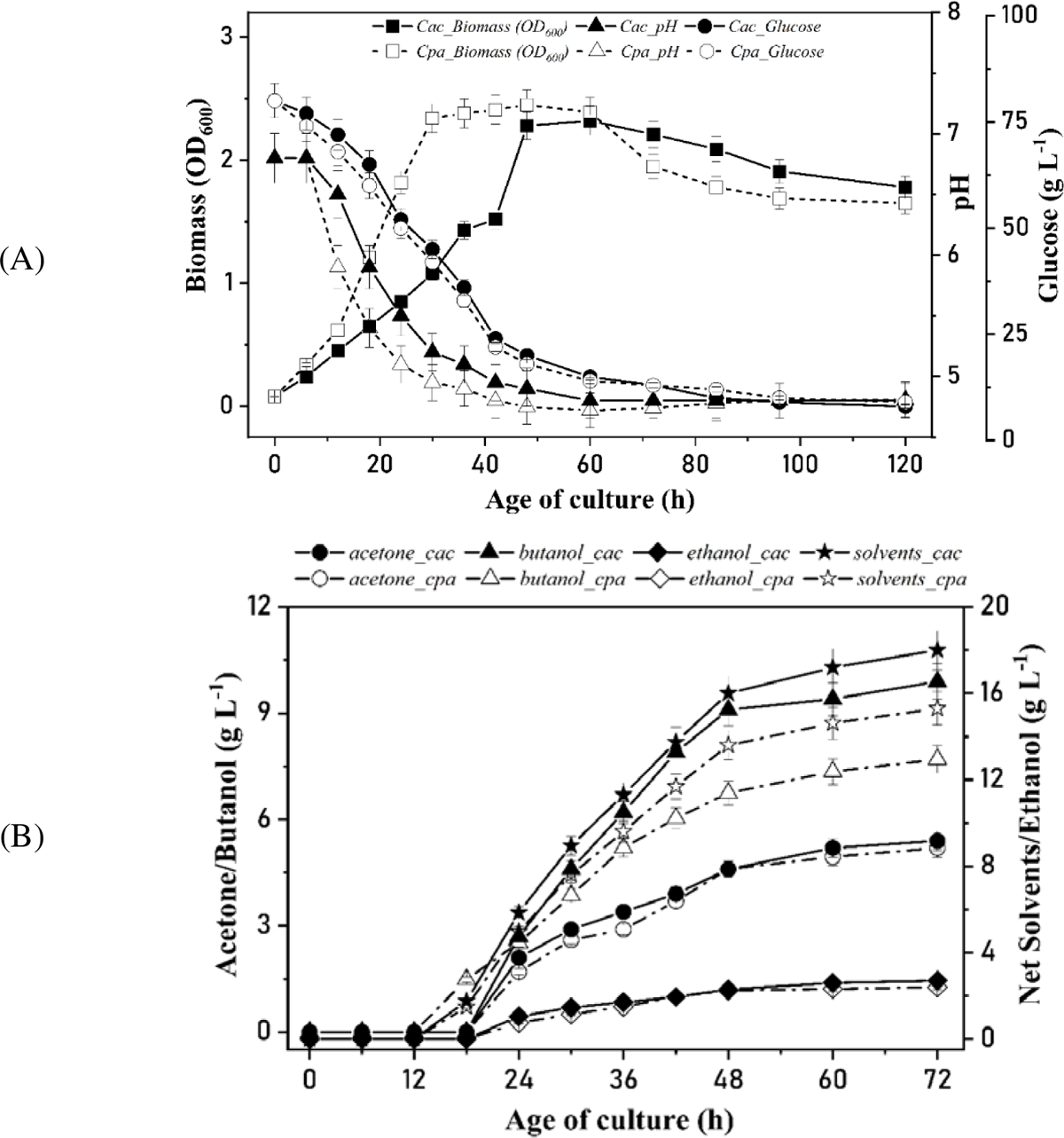
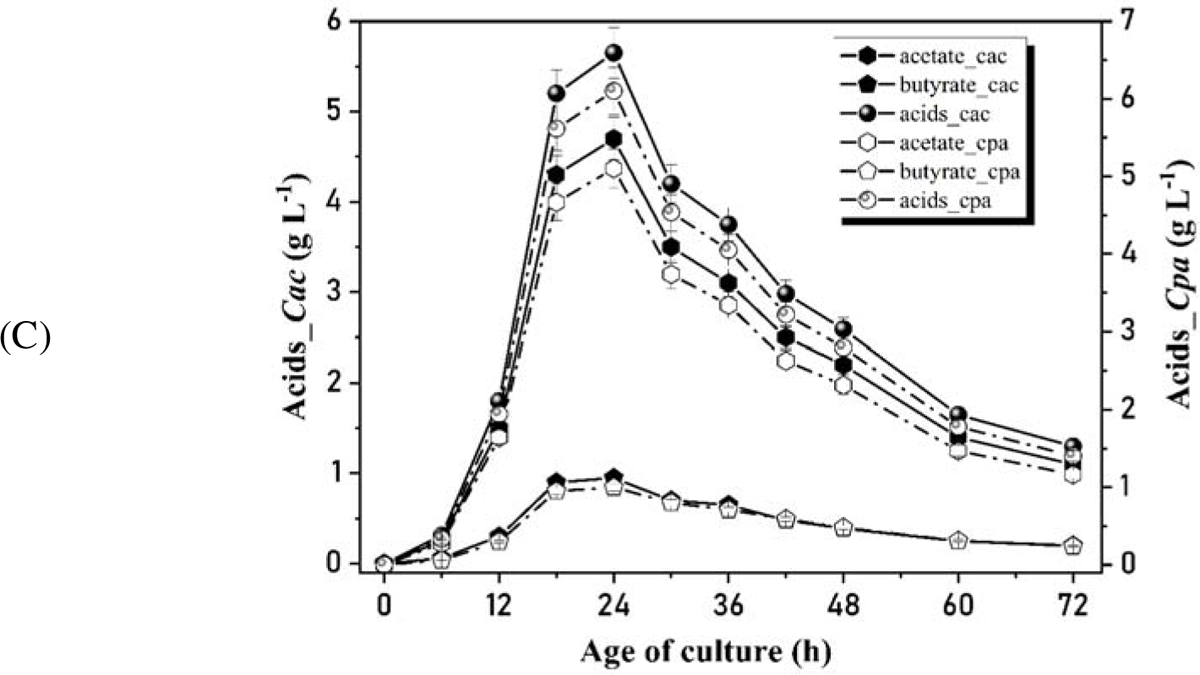
Comparative dynamic profiles for (A) biomass, pH, and glucose consumption; (B) solvents (acetone, ethanol, butanol, and total) production; (C) acids (acetate and butyrate) production.

#### 3.1.3. Comparative growth and butanol production on various carbon sources

In this section, we compare growth and butanol production of *Cac* and *Cpa* on various carbon sources. The aim was to gain insights into their metabolic capabilities and potential for biobutanol production under 15 different carbon sources. Both *Cac* and *Cpa* demonstrated robust growth on C_6_ simple sugars, with slight variations in growth rates observed between strains and carbon substrates. Notably, glucose and fructose supported the highest growth rates for both species, highlighting their preference for these sugars as carbon sources. Qualitative and quantitative results are shown in Table 3 and Figs. 2A & B, respectively.

**Fig. 2.**
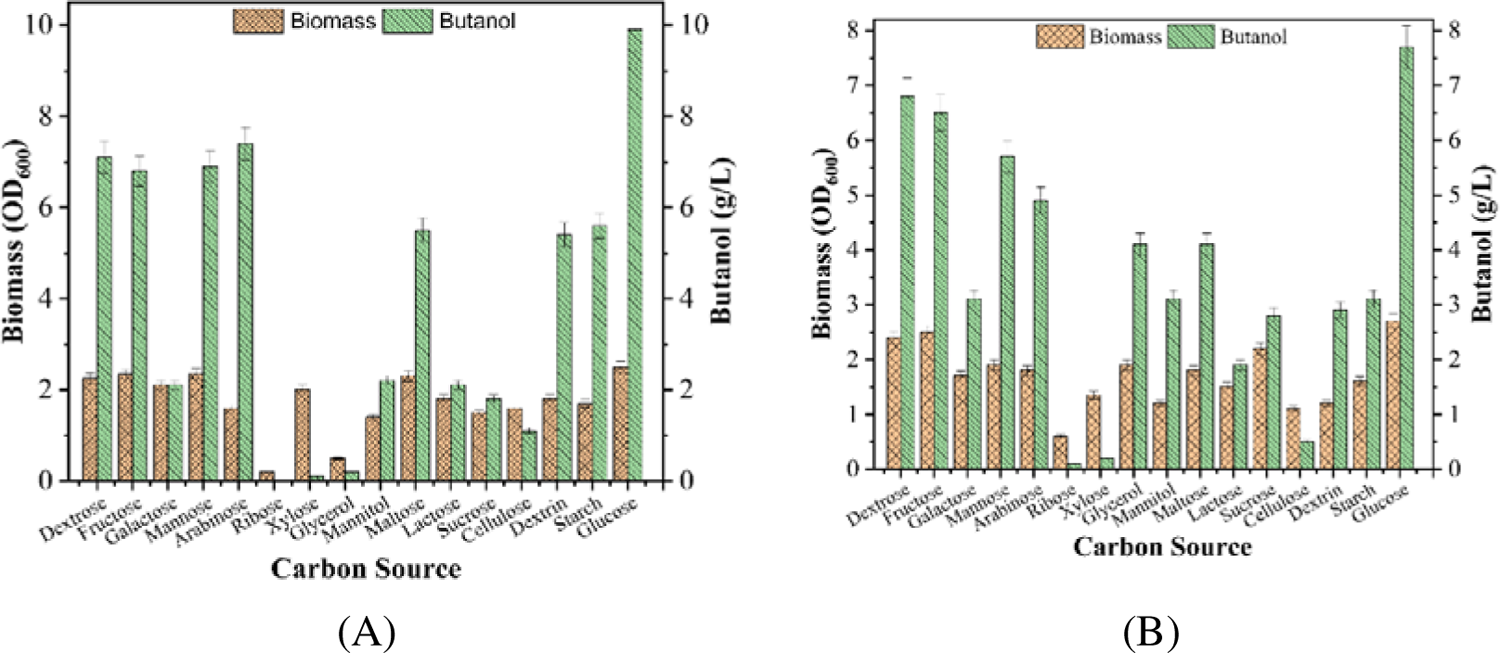
Comparative characterization of (A) *C. acetobutylicum* MTCC 11274 and (B) *C. pasteurianum* MTCC 116 in terms of growth (orange bar) and butanol titer (green bar) under different carbon sources.

**Table 3.** Growth of *Cac* MTCC 11274 and *Cpa* MTCC 116 on various carbon sources.

| Carbon source type | *Cac* MTCC 11274 | *Cpa* MTCC 116 |
| --- | --- | --- |
| **Hexose** |  |  |
| Dextrose | Yes | Yes |
| Fructose | Yes | Yes |
| Galactose | Yes | Yes |
| Mannose | Yes | Yes |
| **Pentose** |  |  |
| Arabinose | Yes | Yes |
| Ribose | No | No |
| Xylose | Yes | Yes (Mild) |
| **Sugar alcohols** |  |  |
| Glycerol | No | Yes |
| Mannitol | Yes | Yes |
| **Disaccharides** |  |  |
| Maltose | Yes | Yes |
| Lactose | Yes | Yes |
| Sucrose | Yes | Yes |
| **Polysaccharide** |  |  |
| Cellulose | Yes (Mild) | Yes (Low) |
| Dextrin | Yes | Yes |
| Starch | Yes | Yes |

Next, we assessed the butanol production capabilities of *Cac* and *Cpa* during fermentation on the same set of carbon substrates. Strikingly, we observed distinct patterns in butanol production between the two species. *Cac* exhibited higher butanol yields on C_6_ sugars, with peak production of 9.9 g L^-1^ (glucose) and 7.7 g L^-1^ (dextrose). *Cac* showed poor butanol production on C_5_ sugars and sugar alcohols. In contrast, *Cpa* displayed superior butanol production on sugar alcohols, reaching a maximum of 4.5 g L^-1^ (glycerol) and 3.8 g L^-1^ (mannitol). These results suggest that the choice of carbon source significantly influences the butanol production capacity of each species.

### 3.2. Optimization of process parameters for co-culture experiments

#### 3.2.1. Butanol maximization as response variable in BBD

The application of coded values from Table 1B to the quadratic regression model for maximizing butanol (i.e., butanol concentration) produced the following equation:

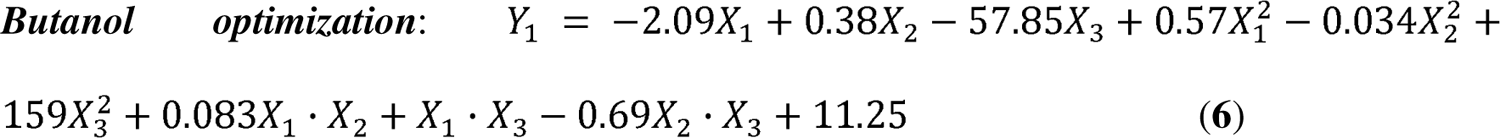

The above expression represents butanol concentration (*Y_1_* in g L^-1^) as a function of inoculum ratio (*X_1_*), NaCl concentration (*X_2_*in g L^-1^), and substrate ratio (*X_3_*).

#### 3.2.2. Biomass maximization as response variable in BBD

Modeling the experimental data for maximizing biomass (i.e., biomass concentration) using the quadratic regression model and the coded values of independent variables listed in Table 1B resulted in the following equation:

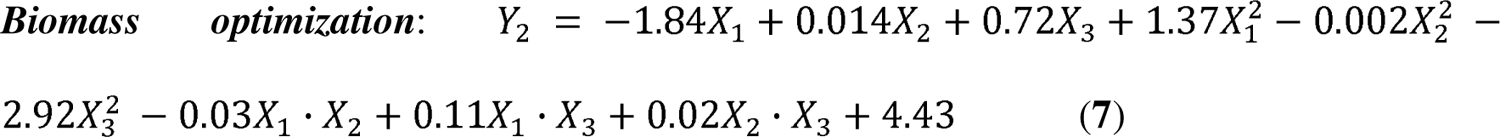

The above expression represents biomass concentration (*Y_2_* in OD_600_) as a function of inoculum ratio (*X_1_*), NaCl concentration (*X_2_*in g L^-1^), and substrate ratio (*X_3_*).

#### 3.2.3. Maximization of ABE concentration as response variable in BBD

Modeling the experimental data to maximize ABE (i.e., ABE concentration) using the quadratic regression model and the coded values of independent variables from Table 1B yielded the following equation:

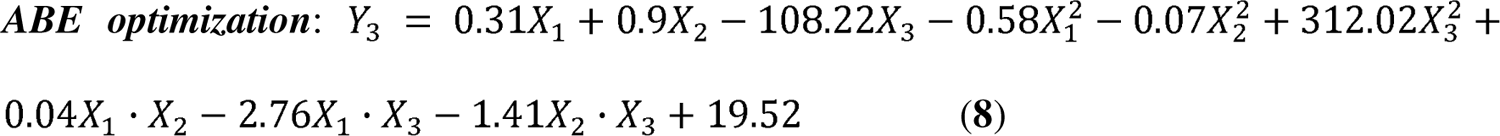

The above expression represents ABE concentration (*Y_3_* in g L^-1^) as a function of inoculum ratio (*X_1_*), NaCl concentration (*X_2_*in g L^-1^), and substrate ratio (*X_3_*).

The comparison between experimental and model-predicted values of the response variable is evident in Table 1B, demonstrating a close agreement and suggesting an optimal fit of the model to the experimental data. Table 4A provides the statistical analysis of the quadratic response model, presenting the coefficients of the models alongside their respective *p*- and *t*-values. Additionally, Table 4B outlines the analysis of variance (ANOVA) results for the quadratic model. ANOVA results indicate the high significance of the regression model (*p* < 0.01), with the lack of fit being non-significant (*p* > 0.05). The coefficient of determination (*R^2^*) exceeding 0.95 in all instances suggests that approximately 95% of the variation in the response variable can be explained by the model. The *R^2^* value surpassing 0.95 further signifies the optimal fit of the model to the experimental data, a conclusion supported by the close agreement between the experimental and model-predicted values of the response variable. The *t*-test, *F*-values, and *p*-values associated with the coefficients of the quadratic model serve as indicators of the relative significance of the corresponding independent variables. A significant *t*-statistic value and a *p*-value less than 0.05 indicate the significance of the coefficient and the corresponding independent variable.

**Table 4.** Results of Box-Behnken Design for optimization of experimental parameters (A) *t*– and *p*– values of the coefficients of quadratic regression models for different response variables.

| Model term | Butanol maximization ($Y_1$) | | Biomass maximization ($Y_2$) | | ABE maximization ($Y_3$) | |
| --- | --- | --- | --- | --- | --- | --- |
| | $t$-value | $p$-value | $t$-value | $p$-value | $t$-value | $p$-value |
| Intercept ($A_0$) | 167.857 | 0.000* | 55.931 | 0.000* | 20.514 | 0.000* |
| *Linear coefficients* |  |  |  |  |  |  |
| Inoculum ratio ($X_1$) | -12.762 | 0.000* | -8.913 | 0.000* | 0.124 | 0.906 |
| NaCl ($X_2$) | -3.118 | 0.026* | 1.338 | 0.238 | 7.132 | 0.001* |
| Substrate ratio ($X_3$) | -88.296 | 0.000* | 2.065 | 0.094 | -25.699 | 0.000* |
| *Square coefficients* |  |  |  |  |  |  |
| Inoculum ration*Inoculum ratio ($X_1^2$) | 2.184 | 0.081* | 9.321 | 0.000* | -0.334 | 0.752 |
| NaCl*NaCl ($X_2^2$) | -26.926 | 0.000* | -2.191 | 0.080 | -8.386 | 0.000* |
| Substrate ratio*Substrate ratio ($X_3^2$) | 112.818 | 0.000* | -3.652 | 0.015* | 32.533 | 0.000* |
| *Interaction coefficients* |  |  |  |  |  |  |
| $X_1 \times X_2$ | 4.737 | 0.005* | -3.005 | 0.030* | 0.314 | 0.767 |
| $X_1 \times X_3$ | 1.715 | 0.147 | 0.326 | 0.758 | -0.699 | 0.516 |
| $X_2 \times X_3$ | -16.987 | 0.000 | 0.796 | 0.462 | -5.101 | 0.004 |

(B) ANOVA for the quadratic regression model
| Source | Butanol maximization ($Y_1$) | | | | Biomass maximization ($Y_2$) | | | | ABE maximization ($Y_3$) | | | |
| --- | --- | --- | --- | --- | --- | --- | --- | --- | --- | --- | --- | --- |
|  | D<br>F | SS | F-value | p-<br>valu<br>e | D<br>F | SS | F-valu<br>e | p-<br>valu<br>e | D<br>F | SS | F-value | p-<br>valu<br>e |
| Regression | 9 | 83.59 | 2477.97 | 0.000* | 9 | 0.248 | 23.13 | 0.001* | 9 | 307.331 | 198.61 | 0.000* |
| Linear | 3 | 29.87 | 2656.29 | 0.000* | 3 | 0.102 | 33.95 | 0.001* | 3 | 99.535 | 263.86 | 0.000* |
| Square | 3 | 52.55 | 4672.97 | 0.000* | 3 | 0.135 | 37.63 | 0.001* | 3 | 203.222 | 393.98 | 0.000* |
| Interaction | 3 | 1.177 | 104.64 | 0.000* | 3 | 0.012 | 3.268 | 0.118 | 3 | 4.574 | 8.879 | 0.019* |
| Residual (Error) | 5 | 0.019 |  |  | 5 | 0.006 |  |  | 5 | 0.860 |  |  |
| Lack of fit | 3 | 0.018 | 46.19 | 0.021 | 3 | 0.006 | * | * | 3 | 0.446 | 0.727 | 0.627 |
| Pure error | 2 | 0.0003 |  |  | 2 | 0.00 |  |  | 26 | 0.414 |  |  |
| Total | 14 | 83.61 |  |  | 14 | 0.254 |  |  | 14 | 308.191 |  |  |

\*Significant $p$ -values, $p \leq 0.05$ , $p$ -value Prob > F
For Butanol maximization: $R^2 = 0.999$ ; Predicted $R^2 = 0.996$ ; Adjusted $R^2 = 0.994$ ; For Biomass maximization: $R^2 = 0.976$ ; Predicted $R^2 = 0.625$ ; Adjusted $R^2 = 0.934$ ; For ABE maximization: $R^2 = 0.997$ ; Predicted $R^2 = 0.974$ ; Adjusted $R^2 = 0.992$

The relative *F*-values of the linear, interaction, and quadratic coefficients provide insights into the significance of the individual effects of independent variables and the extent of interaction among them. According to the ANOVA results presented in Table 4B, the *F*-values for interaction coefficients are considerably smaller than the *F*-value for linear coefficients. This suggests a relatively independent or unrelated effect of individual variables on the concentration of response variables. The *p*-values for all linear and quadratic coefficients are less than 0.05, accompanied by large absolute *t*-values, indicating a significant effect of all variables on the response variables. In contrast, the *t*-values of interaction coefficients are relatively smaller than those of linear and quadratic coefficients, suggesting their comparatively lesser significance. Additionally, the *p*-values for interaction coefficients related to inoculum ratio with NaCl and inoculum ratio with the initial substrate concentration ratio exceed 0.05. This underscores the insignificance of the interaction between these variables, implying that they independently influence the response variable. The relatively small *F*-value for Lack of Fit, accompanied by its associated p-value, indicates that Lack of Fit is not significant when compared to pure error, affirming the overall significance of the model.

The contour plots illustrated in Figs. S4 – S6(A–C) provide a visual representation of the interactions between medium components and their effects on butanol, biomass (OD600), and ABE production. These plots delineate specific regions of the objective content (butanol, biomass, and ABE production) enclosed by contour lines. They offer insights into the interactions among different optimization parameters. e.g., (A) salt and inoculum ratio, (B) substrate ratio and inoculum ratio, (C) substrate ratio and salt. In the given range of optimization parameters (listed in Table 5), Figs. S4D, S5D, & S6D depict the desirability function plots.

**Table 5.** Summary of variables and their optimized values for three objective functions, viz. butanol maximization, biomass maximization, ABE maximization (values in the brackets denotes model predicted maximum values)

| Variable/Response variable | Maximize butanol<br>(11.87 g L<sup>-1</sup>) | Maximize Biomass<br>(OD<sub>600</sub>) (4.06) | Maximize ABE<br>(22.45 g L<sup>-1</sup>) |
| --- | --- | --- | --- |
| X<sub>1</sub> (ml/ml) | 0.3 | 0.3 | 0.46 |
| X<sub>2</sub> (g) | 5.90 | 2.42 | 6.36 |
| X<sub>3</sub> (g L<sup>-1</sup>)/(g L<sup>-1</sup>) | 0 | 0.136 | 0 |

#### 3.2.4. Validation of Experiments at Optimized Parameters

The optimal value of fermentation parameters for all three response variables are summarized in Table 5. The final concentration of butanol, biomass, and ABE was 12.1 ± 0.45, 4.15 ± 0.03, and 23.1 ± 0.55 g L^-1^, respectively, which slightly higher than the results predicted by BBD analysis (i.e., 11.87, 4.06, and 22.45 g L^-1^, respectively). Figs. 3A & B display the dynamic profiles of butanol production and glucose consumption, and the variations in total solvents, total acid production, biomass production, and pH, respectively, during co-culture studies on optimized media to enhance butanol formation (Y_1_).

**Fig. 3.**
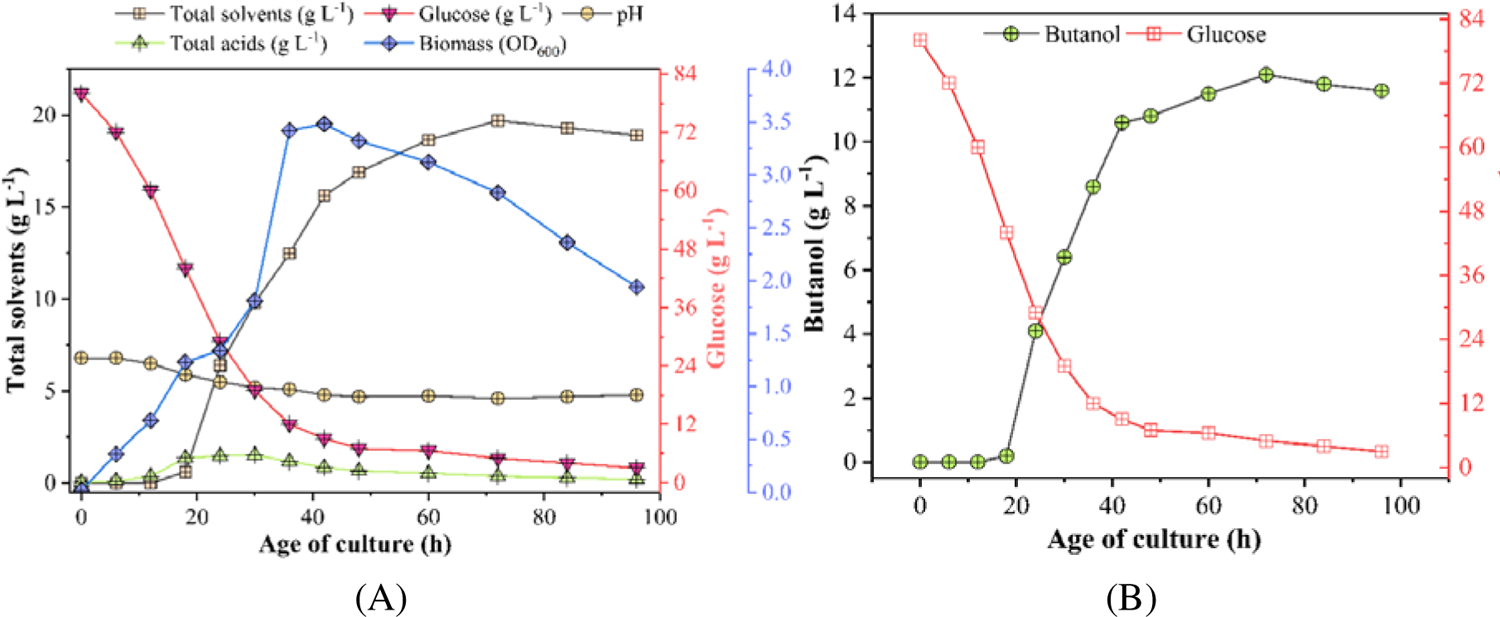
Dynamic profiles of (A) butanol production and glucose consumption, and (B) total solvents, total acid production, biomass production, and pH variation during co-culture studies on optimized media for enhanced butanol formation (Y_1_).

## 4. Conclusions

The insights derived from this study bear substantial significance in the realm of sustainable bioenergy technologies. By unraveling the distinctive physiological attributes of both *C. acetobutylicum* (*Cac*) and *C. pasteurianum* (*Cpa*). Our investigation embarked on a comprehensive journey into the statistical optimization of the co-culture system, comprising *Cac* MTCC 11274 and *Cpa* MTCC 116, with the aim of augmenting biobutanol production from mixed substrates. This endeavor leveraged response surface methodology (RSM) to scrutinize a spectrum of process parameters, including the ratio of *Cac* and *Cpa* inoculum, sodium concentration, and the ratio of xylose to glucose. Upon rigorous analysis, it became evident that salt concentration exerted the most profound influence on biomass, total alcohol, and butanol production. These endeavors culminated in the determination of optimized values for the respective conditions. These optimal conditions yielded highly promising outcomes, with a butanol concentration of 12.1 ± 0.45 g L^-1^ (model prediction: 11.87 g L^-1^), biomass of 4.15 ± 0.03 (model prediction: 4.06 OD_600_), and ABE concentration of 23.1 ± 0.55 g L^-1^ (model prediction: 22.45 g L^-1^). The precision of our models was substantiated through the formulation of quadratic regression models for all three optimization functions.

## Authors’ contributions

**K.K.**: Conceptualization, Methodology, Experiments, Formal Analysis, Validation, Software, Visualization, Investigation, Writing – Original draft, Review and Editing; **S.M.J**: Methodology, Experiments; **L.B.**: Supervision; **V.S. M.:** Supervision, Project administration, Writing – Review and Editing.

## Acknowledgments

Mr. Karan Kumar is grateful to the Prime Minister’s Research Fellowship given by the Ministry of Education, Government of India. Ms. Shraddha acknowledge “Winter INternship in BIOinformatics and Computational Biology-2022 (WINBIOCB – 2022)” organized online at School of Energy Science and Engineering, Indian Institute of Technology Guwahati, Assam, India.

## Compliance with ethical standards

### Ethical approval

This article does not contain any studies with human participants or animals performed by any authors. No conflicts, informed consent, or human or animal rights are applicable to this study.

## Supplementary Information

Supplementary material for this work can be found online with e–version of this paper.

---

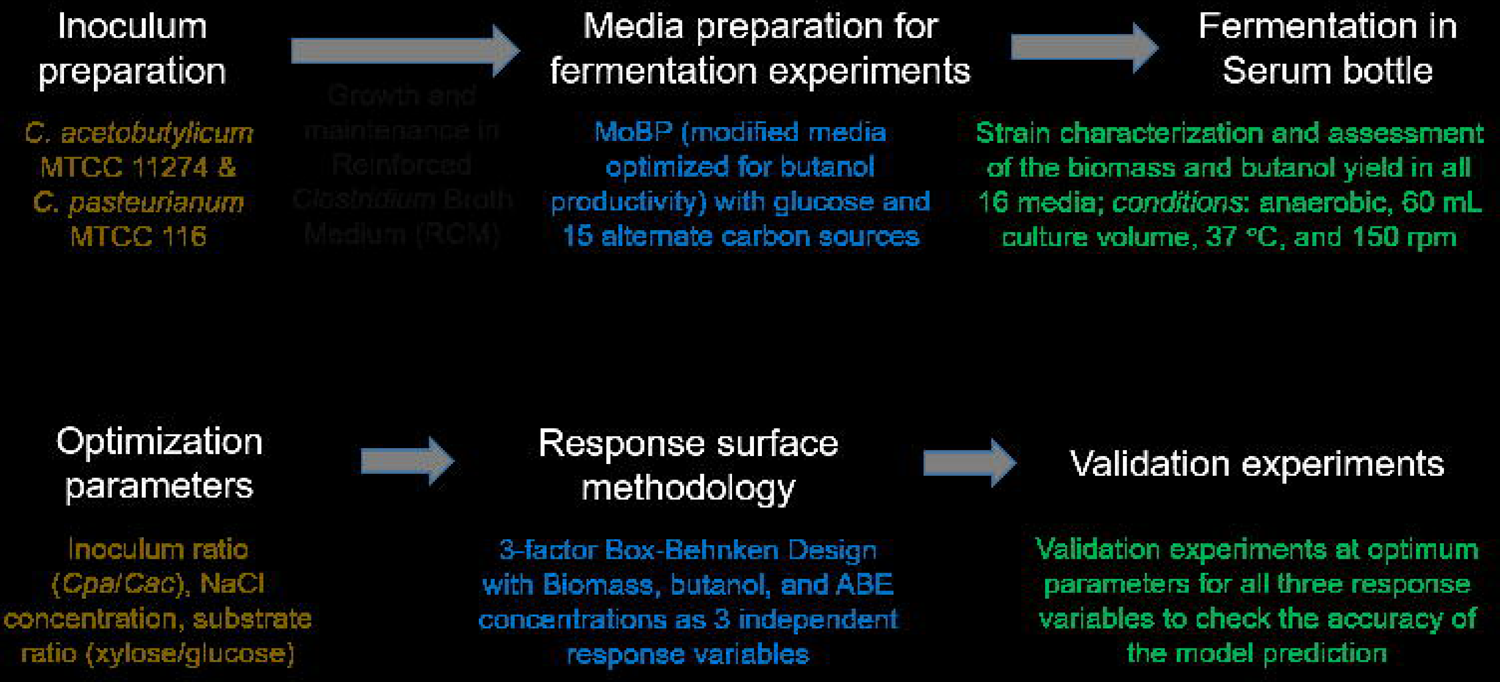

