## Supplemental document for "Acetone-Butanol-Ethanol (ABE) fermentation with Clostridial Co-cultures for Enhanced Biobutanol Production"

**Supplementary file**

**Experimental investigation of ABE fermentation in clostridial co-cultures for consolidated bioprocessing**

Karan Kumar,^1^ Shraddha M. Jadhav,^1^ Lepakshi Barbora,^1^ and Vijayanand S. Moholkar^1,2,*^

^1^ School of Energy Science and Engineering, ^2^ Department of Chemical Engineering, Indian Institute of Technology Guwahati, Guwahati-781039, Assam, India.

| 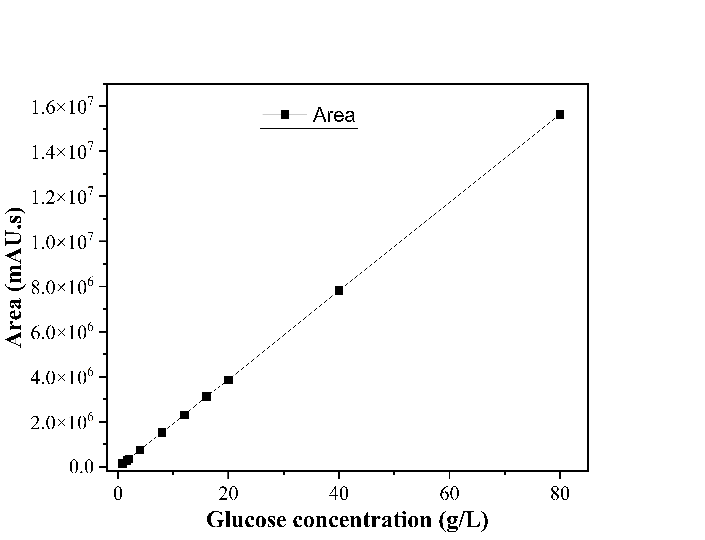 | 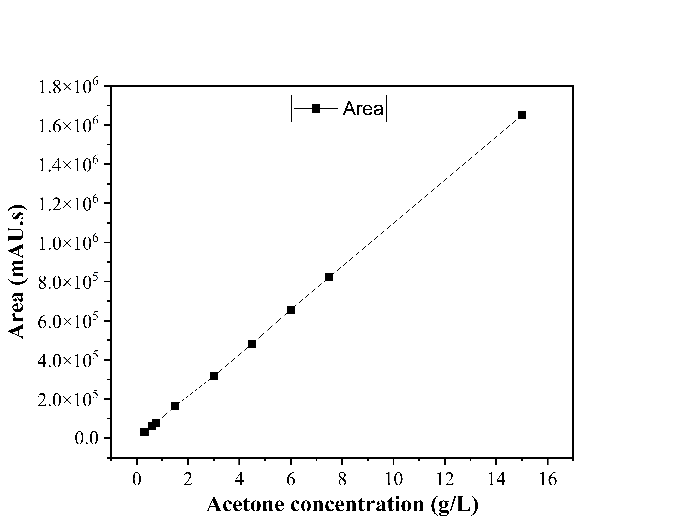 |
| --- | --- |
| 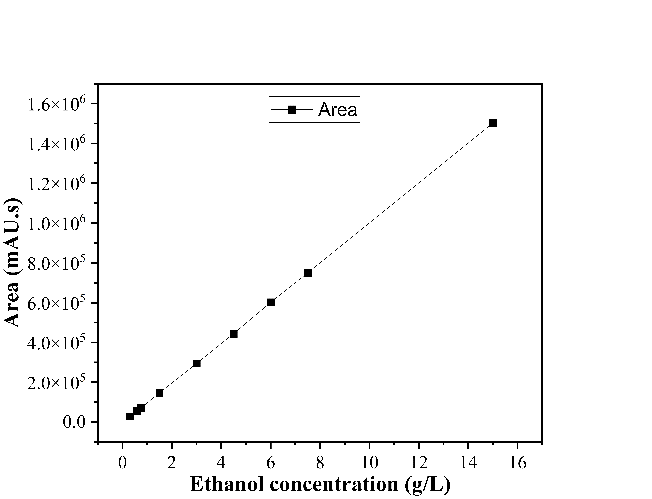 | 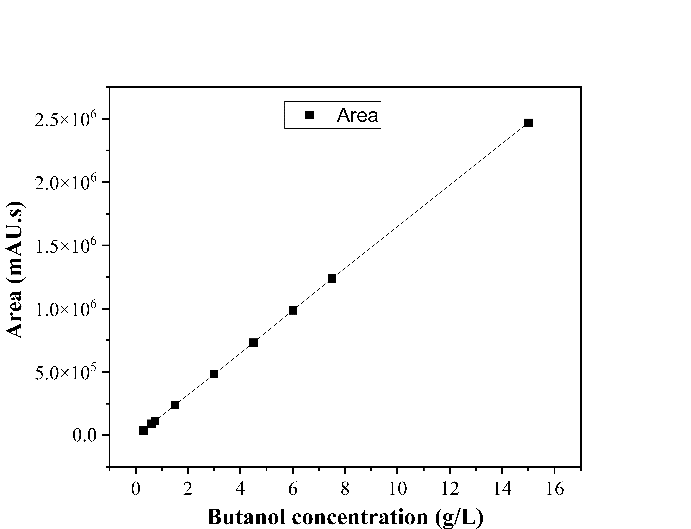 |
| 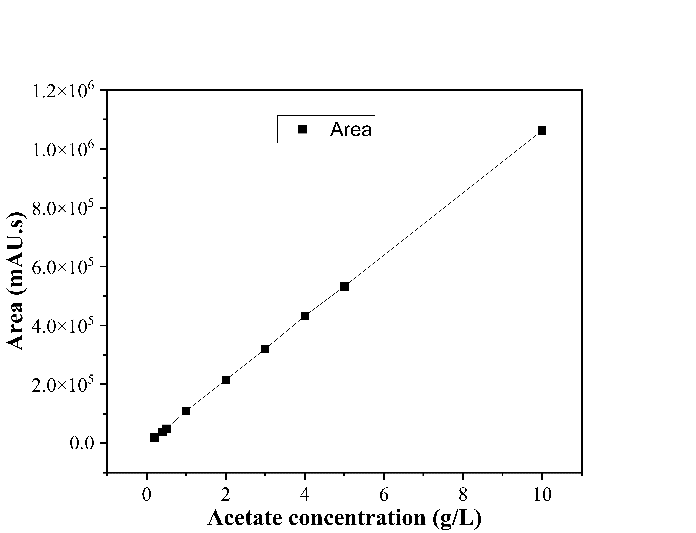 | 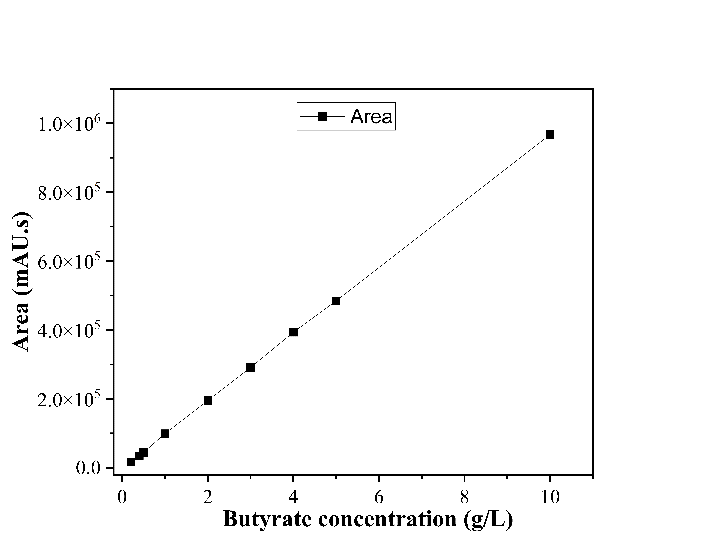 |
| **Figure S1**. Standard calibration plots for glucose, acetone, ethanol, butanol, acetate, and butyrate measured in HPLC | |

| (A) | 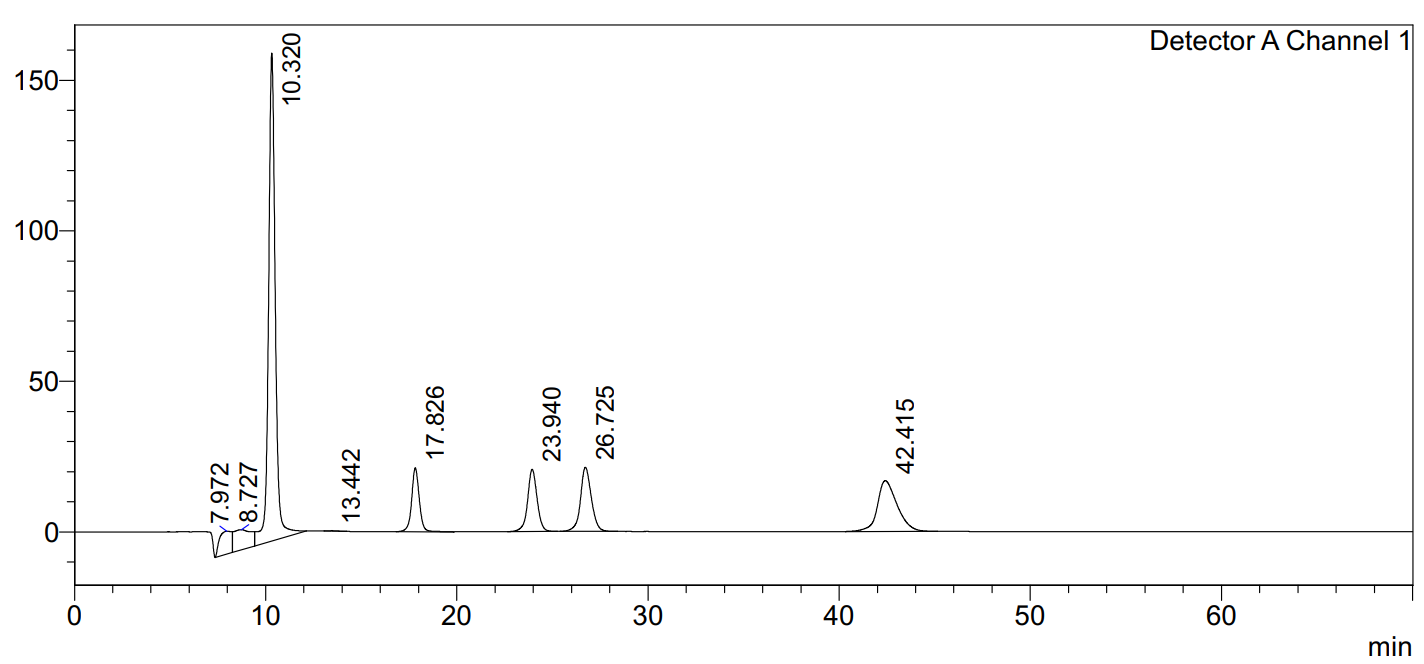 |
| --- | --- |
| (B) | 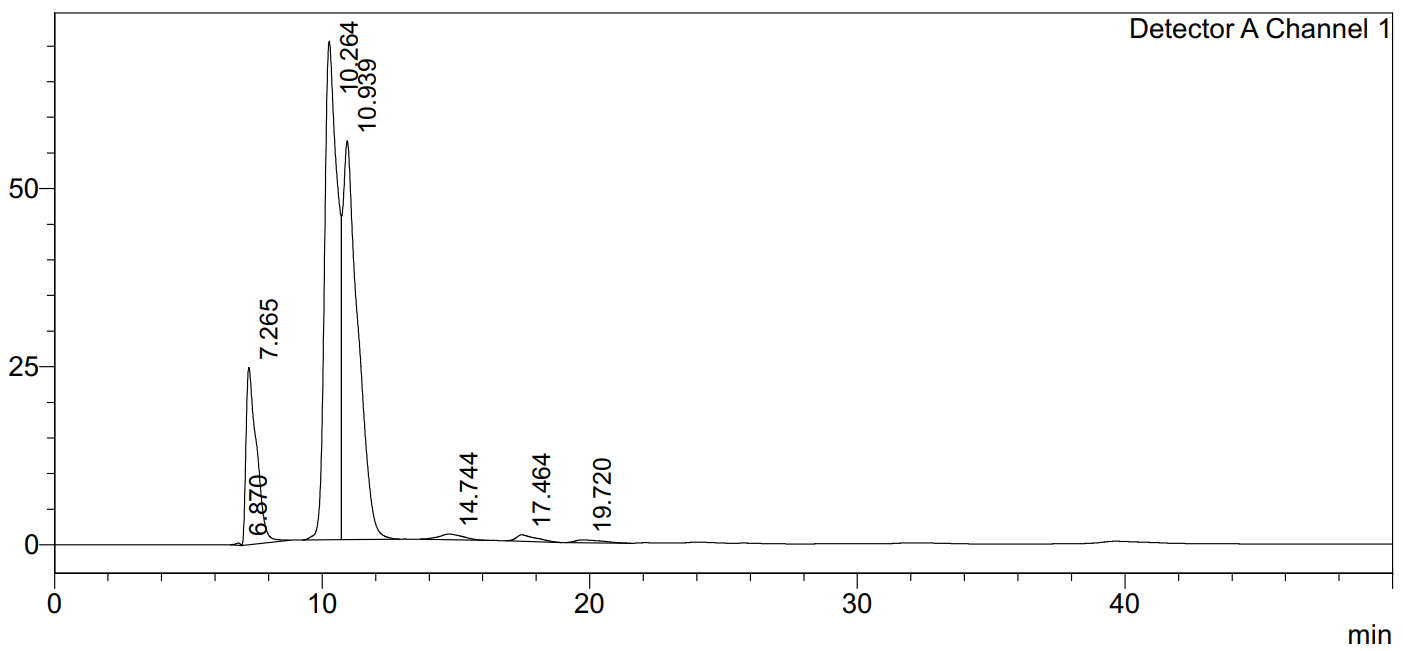 |
| (C) | 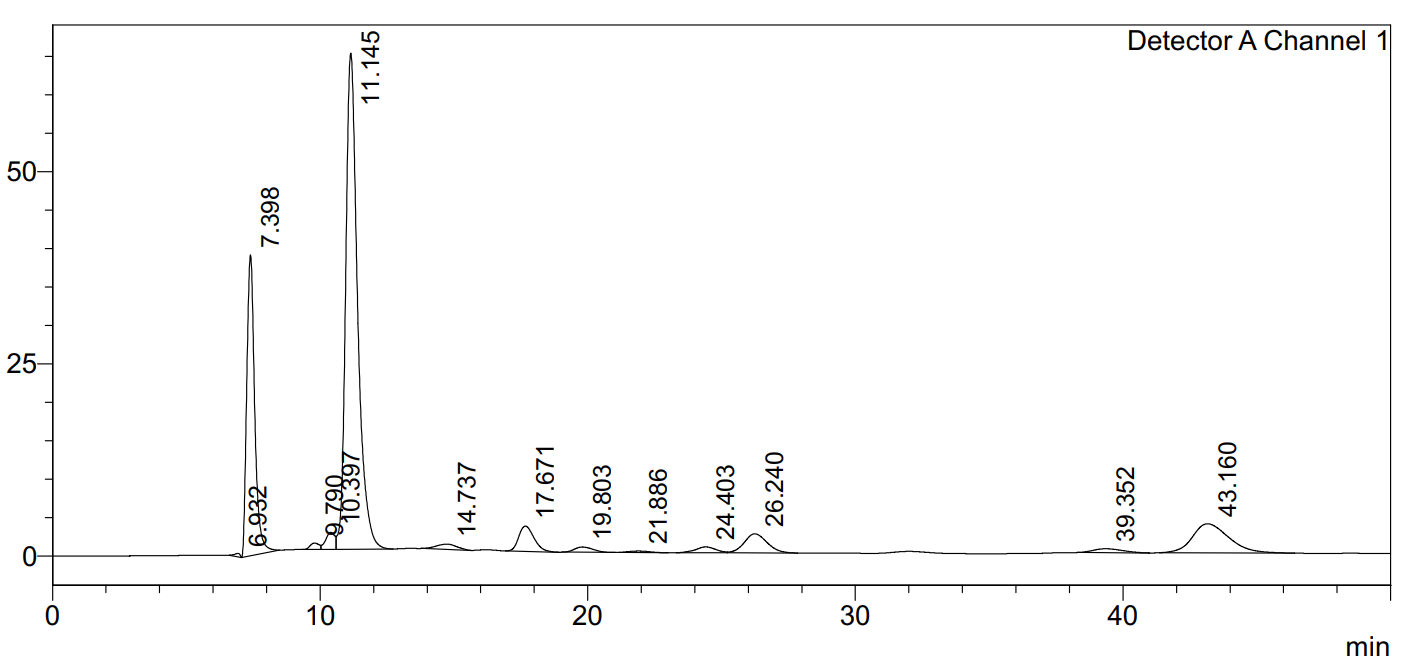 |
| **Figure S2**. HPLC chromatogram showing profiles of compounds (A) standard acetone-butanol-ethanol (ABE) with glucose and acetate, (B) a sample at 0 hours, and (C) a sample at 96 hours of fermentation | |

| 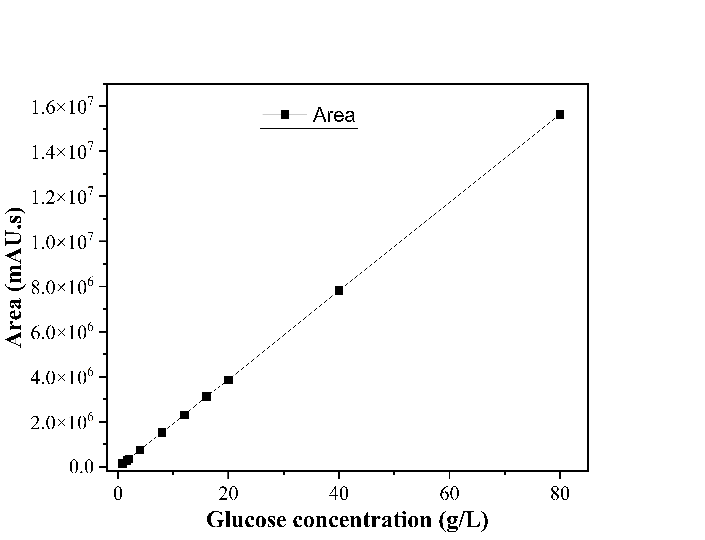 | 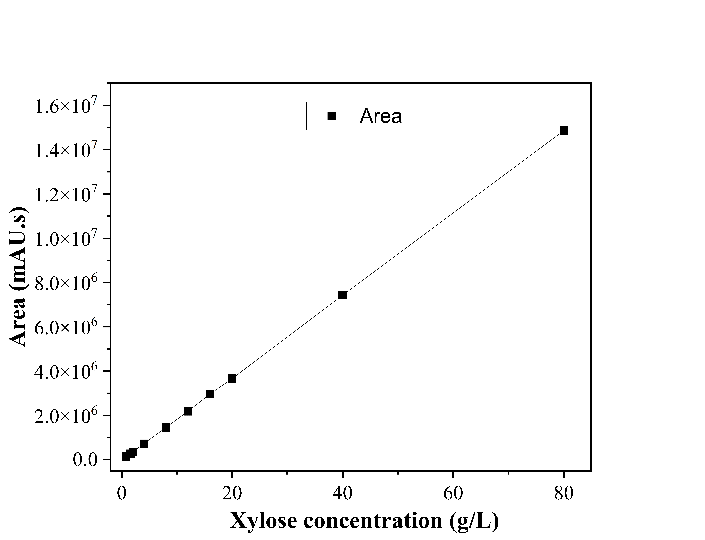 |
| --- | --- |
| **Figure S3**. Standard calibration plots for glucose and xylose measured in HPLC | |

| 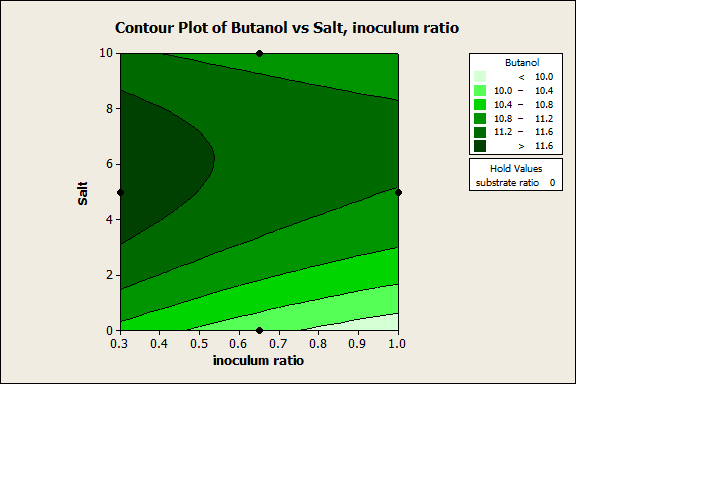  (A) | 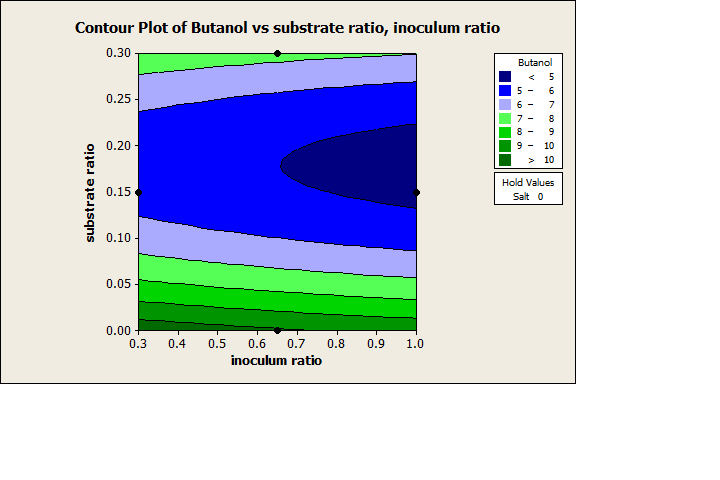  (B) |
| --- | --- |
| 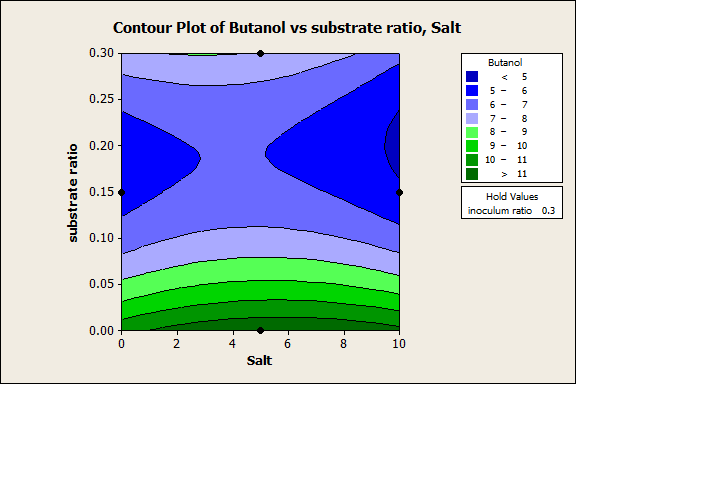  (C) | 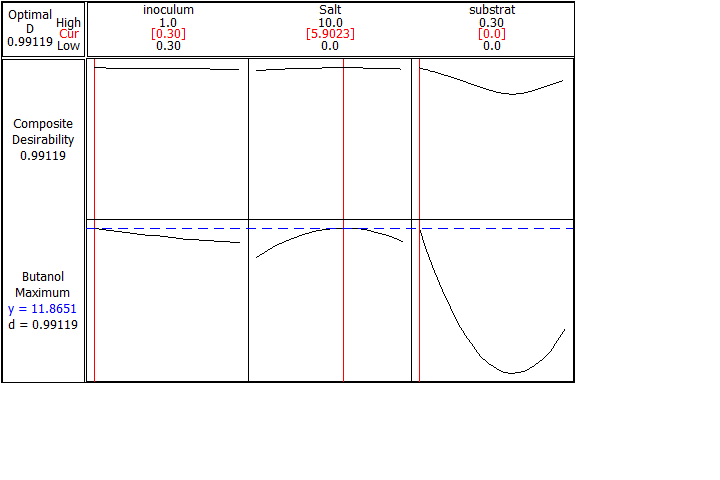  (D) |
| **Figure S4**. Contour plots depicting interactions among different optimization parameters for butanol production (A) salt and inoculum ratio, (B) substrate ratio and inoculum ratio, (C) substrate ratio and salt, and (D) optimization curve | |

| 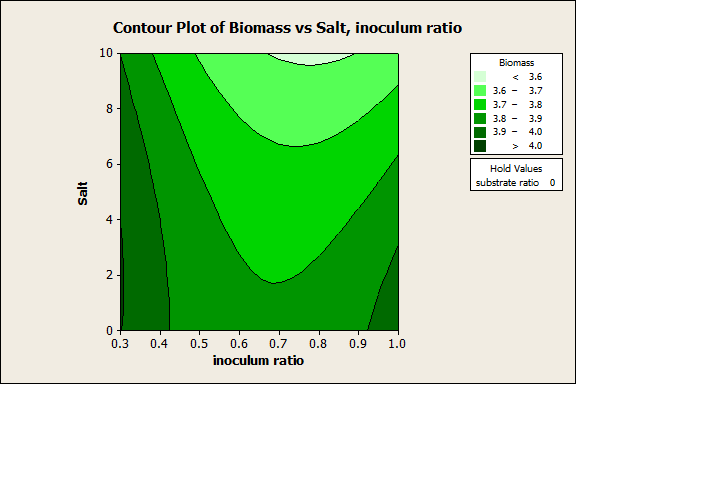  (A) | 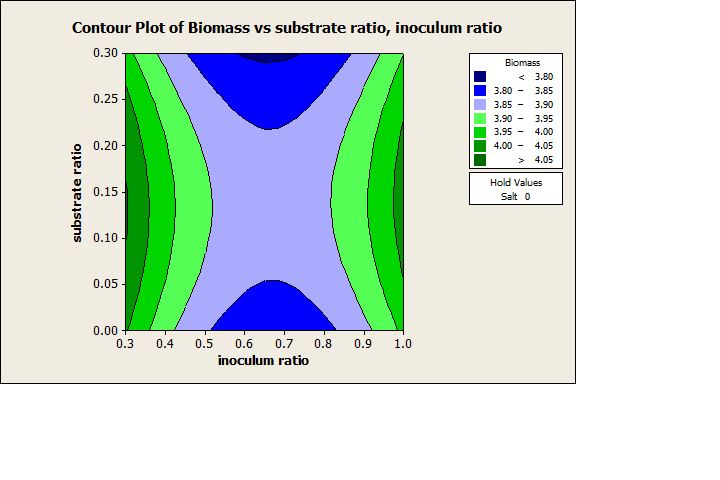  (B) |
| --- | --- |
| 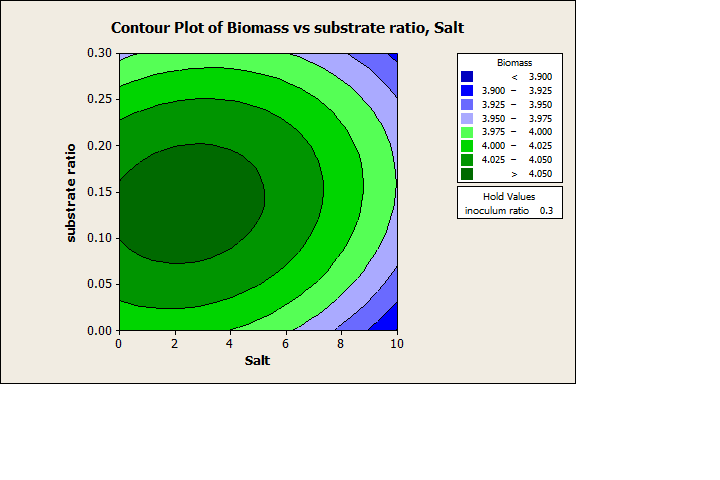  (C) | 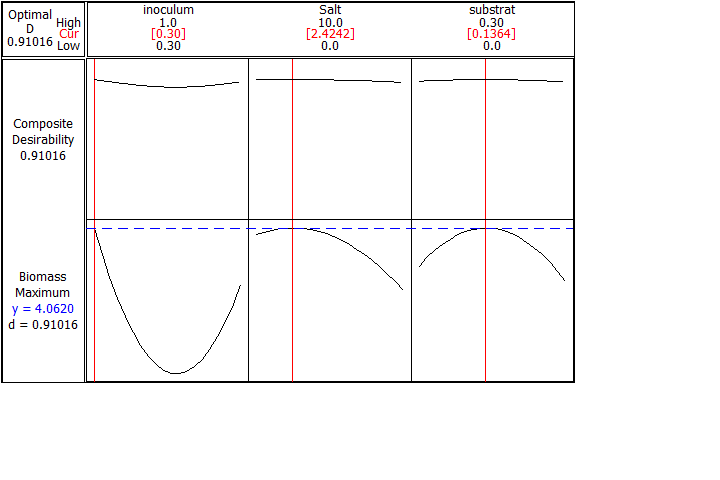  (D) |
| **Figure S5**. Contour plots depicting interactions among different optimization parameters for biomass (OD_600_) production (A) salt and inoculum ratio, (B) substrate ratio and inoculum ratio, (C) substrate ratio and salt, and (D) optimization curve | |

| 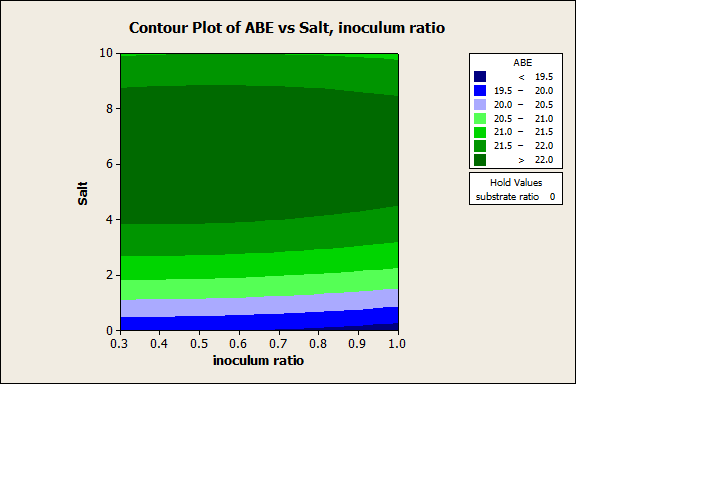  (A) | 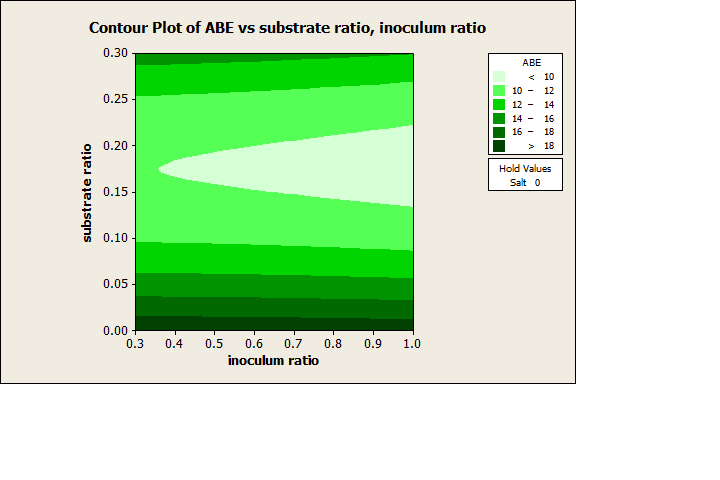  (B) |
| --- | --- |
| 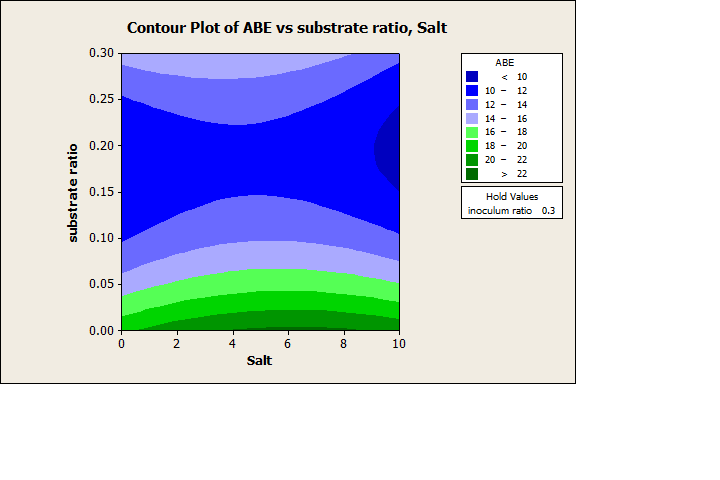  (C) | 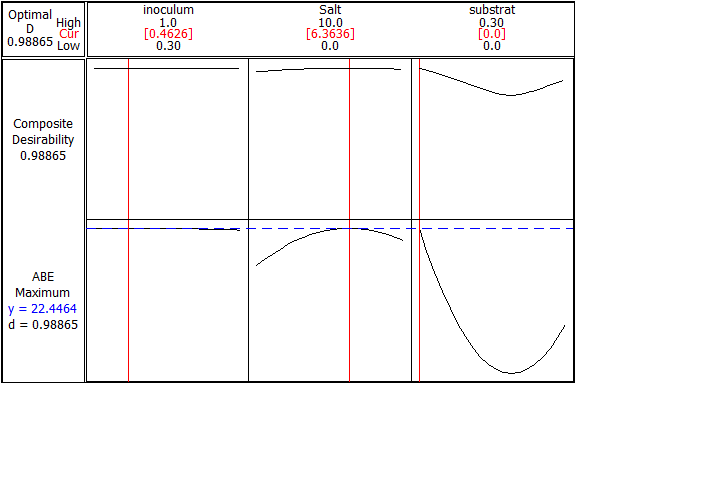  (D) |
| **Figure S6**. Contour plots depicting interactions among different optimization parameters for ABE production (A) salt and inoculum ratio, (B) substrate ratio and inoculum ratio, (C) substrate ratio and salt, and (D) optimization curve | |
